# Primate hippocampus size and organization are predicted by sociality but not diet

**DOI:** 10.1101/713867

**Authors:** Orlin S. Todorov, Vera Weisbecker, Emmanuel Gilissen, Karl Zilles, Alexandra A. de Sousa

## Abstract

The hippocampus is well known for its roles in spatial navigation and memory, but it is organized into regions that have different connections and functional specializations. Notably, the region CA2 has a role in social and not spatial cognition, as is the case for the regions CA1 and CA3 that surround it. Here we investigated the evolution of the hippocampus in terms of its size and organization in relation to the evolution of social and ecological variables in primates, namely home range, diet and different measures of group size. We found that the volumes within the whole cornu ammonis coevolve with group size, while only the volume of CA1 and subiculum can also be predicted by home range. On the other hand, diet, expressed as a shift from folivory toward frugivory, was shown to not be related to hippocampal volume. Interestingly, CA2 was shown to exhibit phylogenetic signal only against certain measures of group size but not with ecological factors. We also found that sex differences in the hippocampus are related to body size sex dimorphism. This is in line with reports of sex differences in hippocampal volume in non-primates that are related to social structure and sex differences in behaviour. Our findings support the notion that in primates, the hippocampus is a mosaic structure evolving in line with social pressures, where certain subsections evolve in line with spatial ability too.

## 1. Introduction

The relationship between behaviour and brain size and proportions has been the topic of intensive research for decades, with works on mammals focusing mainly on the question of how the exceedingly large brains of primates, and particularly humans, could evolve. However, while there is an emerging consensus on the energetic constraints on the evolution of brain enlargement [1, 2], the search for behavioural correlates of relative brain size has a long history of producing a frustrating diversity of results [3]. In primates, there’s a long-standing debate about the degree to which ecological challenges have been met either directly through selection for individuals traits that are adaptations to those ecological challenges, or indirectly through social solutions [4]. Models of primate social intelligence and brain size emphasize social skills, including managing social complexity, theory of mind, social learning, and culture [4, 5]. On the other hand, models of ecological intelligence demonstrate an important impact of home range size and/or diet [6-9] on relative brain size. Evidence about which of these (home range or diet) is the main determinant of brain size is ambiguous [6][7], but both possibly relate to the memory demands of locating and identifying unpredictable food sources or mates, or tool use and social behaviour [10-13]. An important caveat to studies of “intelligence” and brain morphology is the fact that most analyses of brain morphology addressing social and ecological factors across primate evolution only consider how they relate to brain size (absolute or relative). However, there is increasing awareness that more specific aspects of brain organization may better relate to more specific cognitive abilities [14], consistent with long-standing evidence that the brain is a mosaic of different regions, which may respond differentially to selection for specific behaviours [15-18]

The mammalian hippocampus is of particular interest in terms of dissecting the morphological correlates of ecological and social behaviour. It is well known for its roles in both spatial cognition [19] and memory [20], and also has an important role in behavioural inhibition [21] in rodents and primates, including humans. The hippocampus’ role in spatial cognition has been the topic of several comparative analyses related to “ecological intelligence”, and has benefitted from studies in rodents that have revealed a neurophysiological mechanism for mapping spatial coordinates in navigation [19]. The hippocampus contains a population of neurons (‘place cells’) that respond whenever an animal is in a specific location [22] and these produce a dynamic ‘cognitive map’ of the environment by firing in a concerted fashion [23]. Similarly, the entorhinal cortex, a structure neighbouring the hippocampus in the larger “hippocampal complex”, has a population of “grid cells”, which fire when an animal enters an environment with geometrically patterned locations [24]. Another component of hippocampus-related “ecological intelligence” is its essential function in declarative or relational memory possibly through a spatial-based mechanism [25, 26]. The hippocampus also has a role in behavioural inhibition [21] and olfactory memory [27].

While declarative memory is a very broadly relevant cognitive ability it is hard to relate to ecological variables. On the other hand, the hippocampus’ role in spatial cognition is often related to the ecological variable home range, defined as “that part of an animal’s cognitive map of its environment that it chooses to keep updated” [28]. Some studies have suggested a direct link between species’ home range size and species’ hippocampal size. In desert rodents, the bannertail kangaroo rat has relatively low spatial memory requirements and has a small hippocampus, whereas Merriam’s kangaroo rat uses spatial memory to relocate its caches in scattered locations, and larger hippocampus [29]. The “avian hippocampus” in the medial pallial zone is homologous to that in mammals and also functions in spatial memory [30]. This is consistent with the fact that food-storing birds have relatively larger hippocampi [31, 32].

The size and internal organization of the hippocampus is also subject to within-species variation and individual plasticity. Volumetric reorganization of the hippocampus has been related to the occupational specialization in humans [33]. In birds, hippocampal size and structure is plastic, being affected by experience [34], and seasonality [35]. In arboreal primates, a relationship was found between hippocampus size and home range size [36], but overall, this relationship remains unclear [36] [37]. The possibility for a predictive function of the hippocampus is particularly evident from studies of sexual dimorphism in hippocampal size and spatial ability. Whereas male and female meadow voles are sexually dimorphic in their performance on spatial tasks, hippocampus volume, and home range size, pine voles are not [38]. Further, in two other polygamous rodent species the relative size of the hippocampus is greater in males than in females [29], while males and females of the monogamous desert kangaroo rat do not differ in home range nor in spatial ability [39]. Similarly, during breeding season, deer mice are polygynous and males have larger home ranges, and outperform females on spatial tasks [40]. Sex differences in spatial ability and home range size are also related in two species of carnivores - males exhibit larger home ranges and superior spatial ability compared to females in the promiscuous giant pandas, but not in the monogamous Asian small clawed otter [41]. Consistent with the hypothesis that function drives anatomy, the sex differences are reversed in wider ranging females. In a brood parasite bird species, the brown-headed cowbirds, females which travel further than males have larger hippocampi [42] and exhibit superior spatial memory [43].

As of recently, some light has been shed on the role of the hippocampus in social behaviour and cognition. Hippocampal place cells are involved in processing the presence of conspecifics in bats [44] and hippocampal volume has been related to social phobia as part of adjacent circuits in humans [45]. Although the representation system of the hippocampal complex is itself spatial, this coordinate system is capable of processing other spatially representable information – such is the case of its role as a “memory map” for encoding declarative memories [25], or social information [46]. In rats, support for the mechanism comes from studies finding the hippocampus (specifically a substructure described below, CA2) uses place fields to encode information about conspecifics [47]. Given these novel insights into hippocampus function, in species where social behaviour plays an important role, the involvement of the hippocampus in social information processing might be greater. This also has implications for linking social and spatial cognition more generally, as they can be represented in the same cognitive systems [48].

### Hippocampal regions

All fields of the hippocampus formation (retrohippocampus, RH) receive inputs from the entorhinal cortex (EC) along the perforant pathway [49]. Part of it, hippocampus proper, refers to the cornu ammonis (CA) and the fascia dentata (FD); more commonly these same regions are divided up into CA1-3 and the dentate gyrus (DG) (Table 1). DG has traditionally been considered the gateway of the hippocampus because it blocks or filters excitatory afferents from the EC [50]. Sensory and associative projections from the EC synapse in the DG [51]. DG arranges sensory inputs to create a metric spatial representation and is involved in episodic memory and spontaneous exploration of novel environments [52]. DG can be further subdivided into the fascia dentata (FD) and the hilus (part of the CA). Adjacent to the FD, the CA is comprised of four fields arranged in a loop, beginning with the hilus (i.e., CA4) [53]. The hilus is situated along the mossy fiber pathway from the granular stratum of FD to CA3 and is involved in spatial learning and memory retrieval [54]. It has a role in sequence learning [55], and local lesions affect pattern separation, particularly for highly similar inputs [56]. Next are the sequential CA regions in descending order - CA3, CA2, CA1. CA3 receives connections from the mossy fibers of FD, which it projects to CA1 and back, bypassing CA2. There are associational bilateral (ipsilateral and contralateral) connections to CA3 [57]. CA3 can be further divided into subregions: CA3a and CA3b encode spatial information into short-term memory, while CA3c processes environmental geometry along with DG [58]. CA1 receives projections from CA3 and is involved in spatial memory [59]. The spatial properties of CA1 and CA3 are due to these regions being the primary locations of ‘place cells’, responding differentially according to the spatial location of the animal [60]. Adjacent to CA3, the subiculum has inputs from EC and bilateral connections with perirhinal cortex and CA1 [61]. It is a major output of the hippocampus with pronounced dorso-ventral segregation of function: the dorsal component is involved in processing of spatial information and information related to movement and memory, while the ventral is a type of interface between the hippocampus and the hypothalamic–pituitary–adrenal axis, a feedback system that regulates homeostasis and stress [61]. The subiculum receives projections mostly from CA1 and these are organized in a simple pattern - all sections of CA1 project to the subiculum and all parts of the subiculum receive input from CA1 [62]. Moreover, subicular neurons exhibit spatially-selective firing [61] with a robust location signal [63].

**Table 1.**
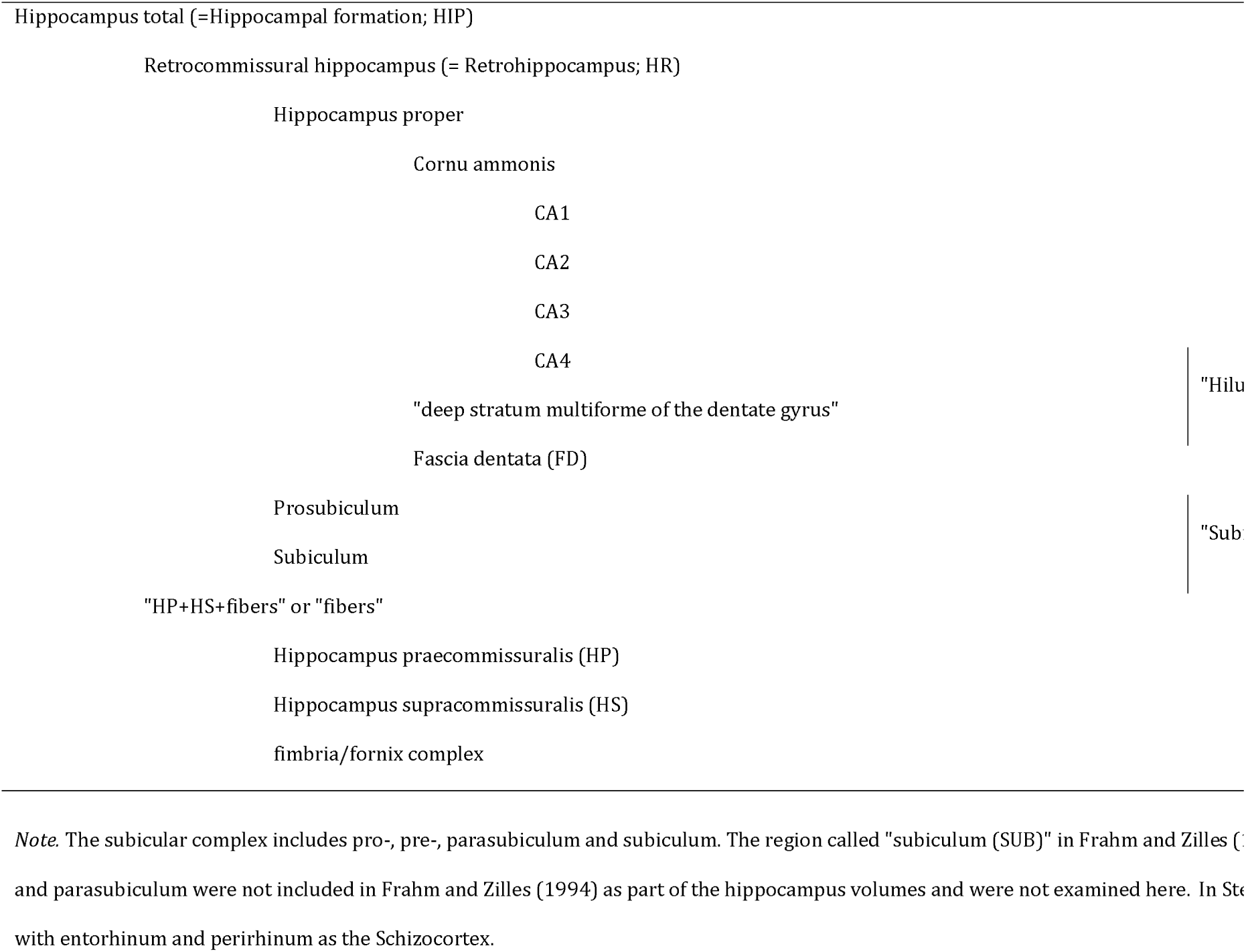
Hippocampal regions investigated.

CA2 has been suggested to act as an interface between emotion and cognition [64]. CA2 receives strong inhibitory inputs from EC, CA3, and DG, and has outputs to CA1 [65]. It is also influenced by many neuromodulators, receiving unique input from hypothalamic nuclei associated with social context, reward, and novelty [64] – supramammillary, paraventricular, median raphe, septal, and the vertical and horizontal limbs of the nucleus of diagonal band of Broca [65]. CA2 has outputs to septum and the supramammillary nucleus. Unlike CA1 and CA3, lesions to CA2 do not affect spatial memory in Morris water maze test, nor impact locomotor ability, anxiety or fear memory in rodents [66]. Rather, CA2 is involved in social memory and recognition of conspecifics [66]. There are some indications its size may be particularly adaptive to social and emotional experiences - decrease in CA2 neuron numbers is associated with schizophrenia and bipolar disorder [67] and stress-related increases in the density of brain-derived neurotrophic factor neurons are greater in CA2 than CA3 [68].

Here we investigate the evolution of hippocampal size and organization in primates, in relation to social and ecological pressures. Given the importance of the hippocampus in spatial cognition, and the subiculum, CA1, CA3, and FD in particular, we predict that these will be related to variation in ecological variables: home range size and/or dietary complexity. Additionally, we predict CA2 volume to be related to social memory, measured through group size. We also expect that amongst brain areas, dimorphism in hippocampal size will be the best predictor of dimorphism in body size.

## 2 Methods

### (a) Anatomical data

The morphometric structure of the hippocampus was determined from previously published volumetric data [69]. For measurements, the retrohippocampus (RH) has been divided into: dentate gyrus (reported in [69] as *fascia dentata*, FD), *hilus* (HIL), *CA3, CA2, CA1*, and *subiculum* (SUB). Volume measurements include the white matter comprising the rest of the hippocampus [69, 70] measured together as *HP+HS+fibers*, that is the hippocampus praecommissuralis (HP) plus the hippocampus supracommissuralis (HS) plus the fibers of the fimbria/fornix complex. Volumes for whole *brain* were taken from the same source [69]. Volumes for *neocortex* (white and grey matter; NEO) were obtained from the same research group [70].

Unpublished data on brain component volumes of males and females were used to determine averages for each sex in a subsample of primates, and correspond to anatomical definitions in [70]. The brain components include 7 telencephalic components: *bulbus olfactorius + bulbus olfactorius accessorius* (bulbus olfactorius accessorius is absent in higher primates; BOL), *lobus piriformis* (palaeocortex and amygdala; PAL), *septum* (septum pellucidum, septum verum, Broca’s diagonal band, bed nuclei of the anterior commissure and stria terminalis; SEP), *striatum* (caudate nucleus, putamen, nucleus accumbens, and the parts of the capsula interna running through the striatum; STR), *schizocortex* (ento- and perirhinal, pre- and parasubicular cortices and the underlying white matter; SCH), *hippocampus* (including all regions; HIP), *neocortex* (white and grey matter; NEO). Included were *diencephalon* (plus globus pallidus without hypophysis; DIE), *mesencephalon* (without substantia reticularis; MES), *cerebellum* (brachium and nuclei pontis, CER), and *medulla oblongata* (plus substantia reticularis; MED). Body weight (BoW) data was available for the same individuals, except for *Miopithecus talapoin* female body weight, which was taken from [71]. Sexual size dimorphism was determined from BoW and calculated as the ratio of male BoW divided by female BoW. Sexual dimorphism in each of the brain structures was calculated as the ratio of the volume in males vs females.

### (b) Social and ecological data

Data were collated from three different sources. Home range area in hectares “HR size average” (HR) were from Powell et al. [7], frugivory “% fruit” were from DeCasien et al. [6], “group size combined” were from [7]. Further, “social group size” data are from [6] and “mean group size” and “mean number of females per group” are from Dunbar et al. [72]. These different studies use different methods for collating the datasets, where it is not always clear whether group size indicates social or foraging group, or whether diet information has been calculated uniformly and reliably.

### (c) Phylogeny

The consensus phylogenetic tree of 43 species of apes and monkeys was obtained from 10k Trees [73] and information about phylogenetic non-independence was incorporated in all analysis. Changes in taxonomic nomenclature were considered for matching species names from the brain dataset to the tree.

### (d) Statistical analysis

All continuous variables were natural log transformed, except for % fruit. Bonferroni correction was applied on the α level (“significance cut-off” of 0.05) on models tested multiple times by dividing it by the number of comparisons with the same dependent variable (4 models with different group size measures resulting in corrected α of 0.0125). Analyses were run on R version 3.6.1 [74] using the packages phytools [75] and caper [76]. Using the fastanc function in phytools we estimated the ancestral states and painted them on the tree using Fancytree. We used caper for all PGLS analyses. Phylogenetic signal (Pagel’s λ) was estimated using Maximum Likelihood and kappa (k) and delta (δ) were fixed to 1. We tested four ‘full’ models including home range size, fraction fruit and each of four different measures of group size against hippocampus and hippocampal region volumes. Additionally, we explored the relationship between neocortex and brain volume with hippocampus volume. Means square statistics were obtained via sequential sum of squares ANOVA.

The volume of the region of interest was always used as the dependent variable in our models, and brain volume was included as a covariate. All variables were shown to be normally distributed, and variance inflation factors of each models were shown to be <3.5 meaning that there was no problem with collinearity. Interactions between predictors were not included as to avoid high cross-collinearity.

We also tested additional single variable models including either only home range, diet or group sizes against hippocampal regional volumes, also correcting for total brain volume. This was done because the ‘full’ model resulted in sample sizes between 20 and 30, while running the separate models mostly utilised the full dataset of 43 species. For results of these models see the Supplementary material.

Additionally, all four ‘full’ models were evaluated and ranked using AIC (Akaike Information Criterion). [77].

All data (including anatomical, social and ecological variables), code, phylogenetic trees and analysis outputs are included in the supplementary material.

## 3. Results

### (a) Ancestral state estimation

An exploratory ancestral state estimation revealed that in species where relative hippocampal volume has decreased (calculated as the residuals from the phylogenetic regression with total brain volume) have nonetheless undergone an increase in absolute hippocampus volume (Fig. 2). We further tested this observation using PGLS and found that hippocampus volume increased with a shallower slope compared to both brain and neocortex volumes i.e. species that evolve towards greater neocorticalization have smaller relative hippocampi. (See Neocortex section). An exception is the pygmy marmoset (*Callithrix pygmaea)*, for which both absolute and relative volume have decreased. This finding is unsurprising due to the expected effects of dwarfism in this species and the limitation this exerts on brain size [78]. In the case of the lar gibbon (*Hylobates lar)* the analysis revealed an increase in both volumes from the ancestral state, possibly reflecting the complexity of its habitats and the subsequent expansion of both hippocampus volume and brain volume.

**Figure 1.**
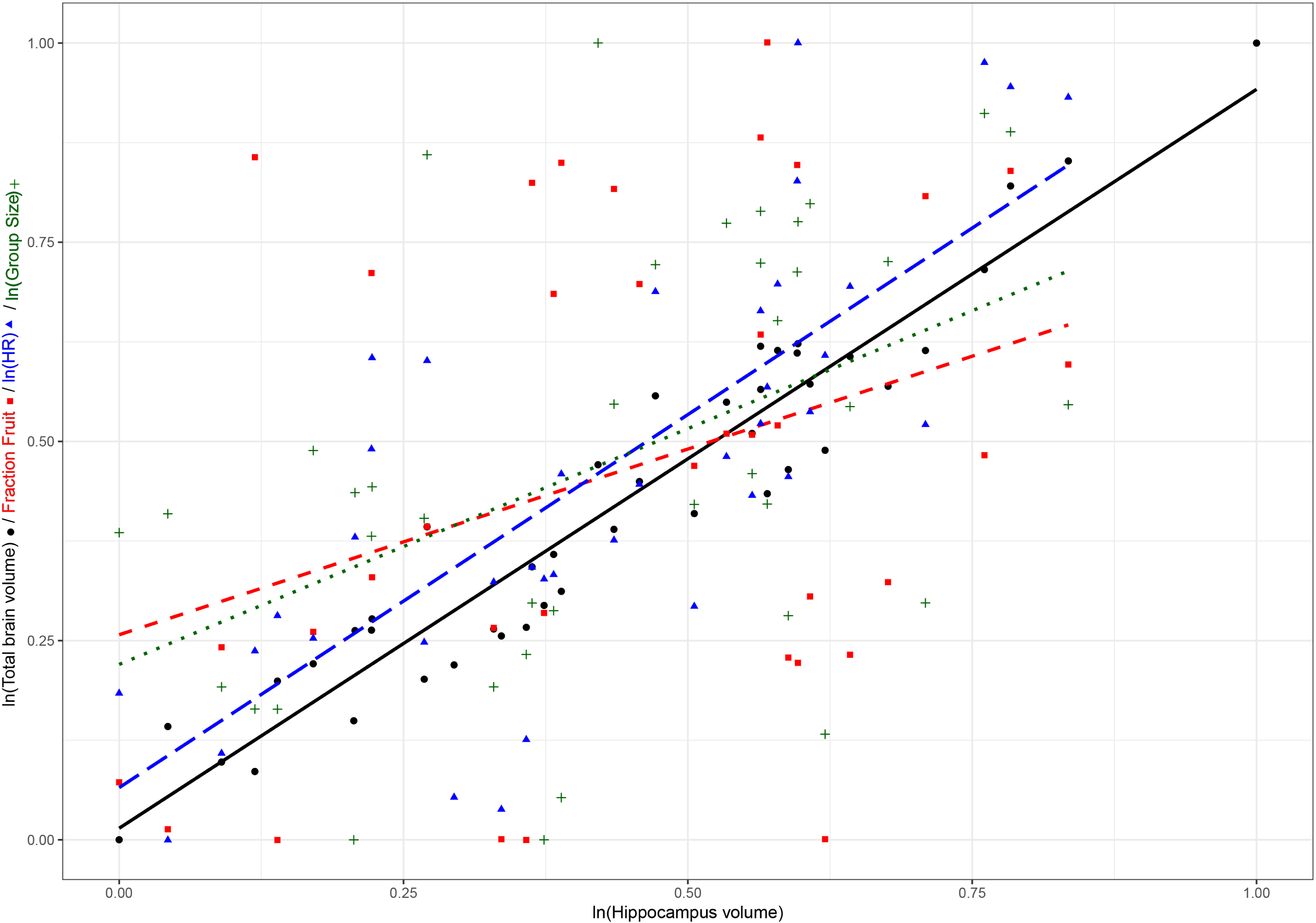
Hippocampus of *Miopithecus talapoin*. LV – Lateral ventricle, FD – Fascia dentata.

**Figure 2.**
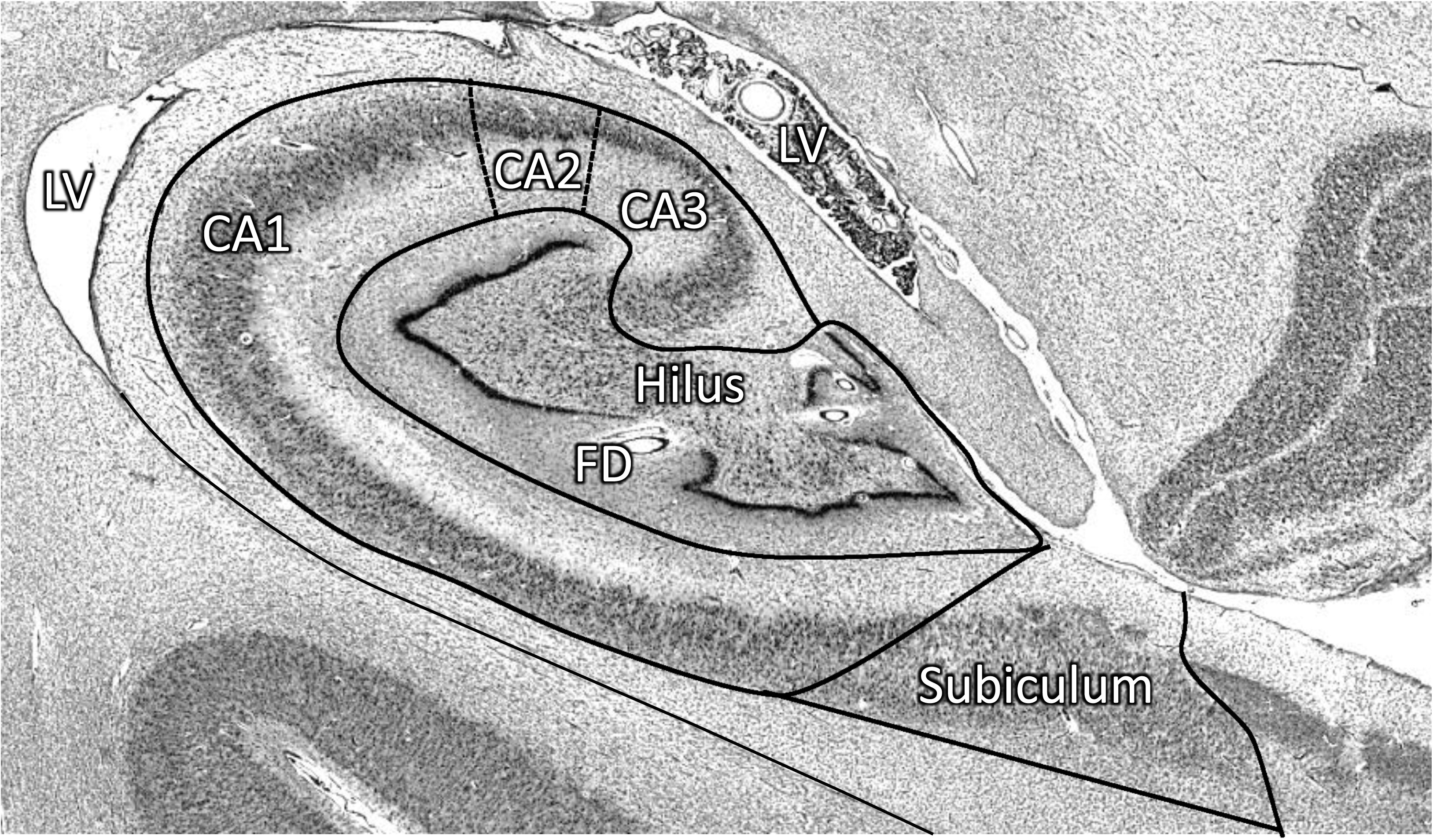
Ancestral state estimations of absolute hippocampal volume (left), and the residuals from the phylogenetic regression with total brain size (right). We observe that most species that had increase in absolute hippocampal volume had a reciprocal decrease in hippocampal volume relative to the whole brain.

**Figure 3.**
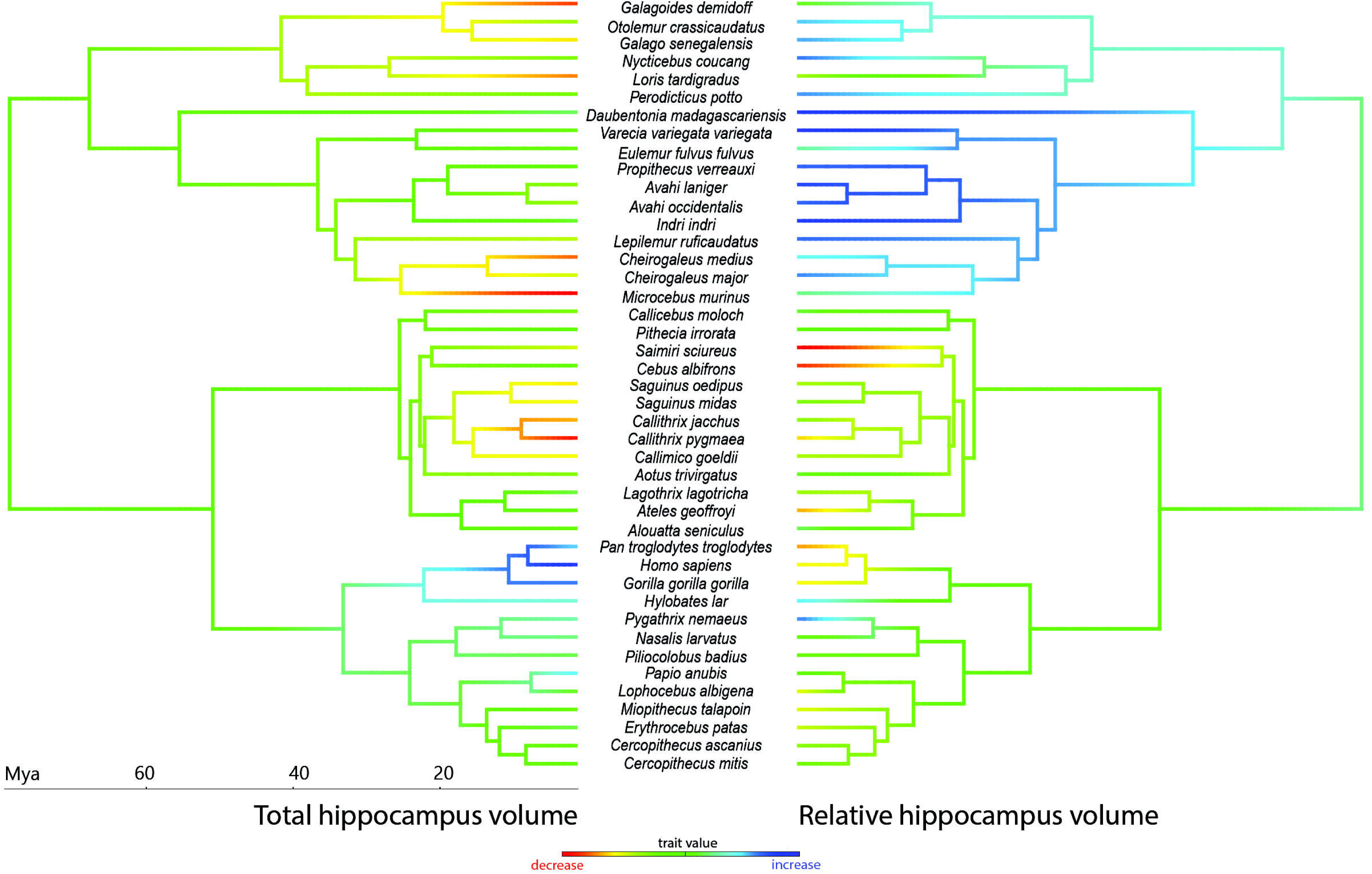
Plot of regression of total brain volume (black solid line and black circles), fraction fruit (red small-dashed line and red squares) and home range (blue long-dashed line and blue triangles), and group size (green dotted line and green pluses) against hippocampal volume.

### (b) PGLS

Testing the ‘full’ models with all four different group size measures separately yielded comparable results. Shown in Table 2 are the results with the groups size measure resulting in the largest sample size – group size from Dunbar [72]. Hippocampus and all regional volumes besides subiculum and hilus could be predicted by group size, home range was shown to be a significant predictor of subiculum and CA1, while fraction fruit was not significantly related to any of the hippocampal structures. The results of the other three models are included in the supplementary material.

**Table 2.** ANOVA output from testing the full model (with Group size from Dunbar) versus hippocampal and regional volumes. On Shown are means squares from the sequential SS ANOVA, p-values and lambda values of the phylogenetic signal of the residuals.

|  | Hippocampus |  | HP+HS+ fibers |  | Retrohippocampus |  | Subiculum |  | Hilus |  | CA1 |  |
| --- | --- | --- | --- | --- | --- | --- | --- | --- | --- | --- | --- | --- |
|  | Mean sq | p | Mean sq | p | Mean sq | p | Mean sq | p | Mean sq | p | Mean sq | p |
| Total Brain | 0.397 | <0.001* | 0.568 | <0.001* | 0.354 | <0.001* | 0.342 | <0.001* | 0.441 | <0.001* | 0.367 | <0.001* |
| Home Range | 0.002 | 0.074 | 0 | 0.58 | 0.005 | 0.02 | 0.014 | 0.007* | 0.009 | 0.04 | 0.012 | 0.001 |
| Group Size (Dunbar) | 0.012 | <0.001* | 0.011 | 0.002* | 0.013 | <0.001* | 0.008 | 0.04 | 0.007 | 0.06 | 0.013 | <0.00 |
| Fraction Fruit | 0.001 | 0.25 | 0 | 0.94 | 0.001 | 0.27 | 0 | 0.74 | 0.002 | 0.38 | 0 | 0.34 |
| Residuals | 0 |  | 0.001 |  | 0 |  | 0.001 |  | 0.002 |  | 0 |  |
| $\lambda$ | 0.59 | | 0.56 | | 0.59 | | 0.47 | | 0.22 | | 0.58 | |

Additionally, each of the four models using different group size measures were compared using AIC (Table 3) and while female group size (from Dunbar [72]) was shown to produce best fitting models in most cases, the sample size was the lowest (N=20) eliminating more than half of the species included in the dataset. In order to utilise our full dataset of 43 species, we also ran separate models including only 1 class of predictors (ecological, social or dietary). The results were concordant with the ‘full’ models and are included in the supplementary material.

**Table 3.** Model fit ranking of all four group size measures. Displayed are the AIC scores and all m are bolded.

|  | Group size (Powell) | Social group size (DeCasien) | Group size (Dunbar) | Female group (Dunbar) |
| --- | --- | --- | --- | --- |
| Hippocampus | -3.7 | **-6.8** | 9.0 | **-6.7** |
| HP+HS+ fibers | 10.3 | 9.5 | **4.2** | **6.1** |
| Retrohippocampus | 0.2 | **-1.8** | **-2.2** | **-2.5** |
| Subiculum | 25.5 | 23.1 | 24.3 | **17.1** |
| Hilus | 33.9 | 34.9 | 34.1 | **12.6** |
| CA1 | 4.3 | **2.5** | **1.5** | 5.4 |
| CA2 | 4.9 | **2.1** | **1.4** | **2** |
| CA3 | -14 | **-4.9** | -9.7 | -7 |
| FD | 7.7 | 6.6 | 6.9 | **-4.2** |

### (d) Neocortex

Following up on the observation that 1) both hippocampus and all its subcomponents were positively related to brain volume, 2) many interactions between predictors and brain volume were yielding negative slopes and 3) with increase in absolute hippocampal volume in some species there was nonetheless a decrease in the relative hippocampal volume, we investigated whether that relationship is driven by variation in neocortex volume as it comprises significant proportion of the total brain volume. We found that hippocampal volume is strongly negatively related to neocortex volume (λ = 0, slope = −3.81, t= −8.11, p<0.0001, 3, 40 df) even after accounting for brain volume (see Supplementary results).

### (e) Sexual size dimorphism

We further explored the relationship between somatic and brain structure sexual dimorphism in a separate dataset of 12 primate species. Somatic sexual dimorphism was best predicted by hippocampus volume dimorphism, (λ = 0, slope = 1.87, Std. error = 0.35, t = 5.19, p=0.0004 on 1 and 10 df). Even though dimorphism in mesencephalon (λ = 0, slope = 1.35, Std. error = 0.45, t = 2.96, p=0.014 on 1 and 10 df) and lobus piriformis (λ = 0.78, slope = 0.67, Std. error = 0.23, t = 2.83, p=0.018 on 1 and 10 df) were also significant predictors of somatic sexual dimorphism, these relationships didn’t stand after correction for multiple comparisons. The new level of α for this batch of analysis was fixed to 0.0045 (dividing 0.05 by 11 structures) and was sufficed by hippocampus volume alone. None of the other structure volumes (OBL, CER, DIE, BOL, SCH, SEP, STR, NEO) showed a relationship with somatic sexual dimorphism.

## 3. Discussion

We find that in primates, hippocampal volume and most of its subcomponents can be reliably predicted by different measures of group size and home range to a certain extent, but not diet. Moreover, we suggest that as brains get larger, the neocortex may take on functions shared with the hippocampus and thus hippocampus size relative to the rest of the brain gets smaller. Alternatively, the size of the hippocampus might be under strong developmental constraint. Hippocampal structures crucial to spatial memory, CA1 and subiculum, evolve in line with ecological (spatial) and social demands. CA2, CA3 and fascia dentata were shown to evolve in line only with social demands, unlike the hilus, for which volume could not be predicted by any of our models. No relationship between hippocampal volume and any of its subcomponents was detected with increased fruit consumption in the primate’s diet. First, neocorticalization outpaces the enlargement of the hippocampus, as indicated in the ancestral state estimation and the subsequent follow-up analysis. This is likely due to a reallocation of functions such as memory, spatial cognition, and inhibition from the hippocampus to the neocortex. With neocorticalization, parallel systems are thought to have emerged, leading to an increased neocortex ratio [79] and allocation of functions to the neocortex [80]. Whereas in smaller brained species the hippocampus is of utmost importance in many cognitive abilities, as the neocortex expands there may be a greater proportion of these functions allocated to it, or the neocortex might be taking up on an array of new social functions that do not exist in smaller brained species. The neocortex, like the hippocampus, provides mappings used in information acquisition, retention and use. Compared to rodents, in highly neocorticalized humans, the hippocampus may not have as prominent a role in spatial cognition (especially when compare to its well-known role in human memory) [81]. On the flip side, in primates, the neocortex may also have an increased role in spatial processing. Parietal association areas of the neocortex are also crucial to spatial perception and may provide navigational information and are the focus of spatial cognition studies [82]. The interplay between the parietal and hippocampal neural networks remains poorly understood [83] although it has been suggested that both are involved in spatial navigation. Parietal representations provide an egocentric frame of reference and may map movements along a route according to route-centred positional information [84].

Second, of the hippocampal regions, both CA1 and CA3 residuals show phylogenetic signal and coevolve with home range (CA1) and group size (CA1 and CA3) when we test single variable models (see Supplement for data on phylogenetic signal within each separate model). This is consistent with the notion that the hippocampus is involved in both social and ecological behaviour [44, 47]. Compared to other brain component volumes, hippocampus volume was found to be the best predictor for cognitive tasks measuring executive function in primates [85]. This is the first study linking theses specific hippocampal substructures to both social and ecological factors across primates. This is in line with work in other taxa linking species-specific requirements for spatial memory and hippocampus volume [29], but the implication - which would benefit from future study - is that in primates the role of the hippocampus may be even more influenced by social factors.

We found no relationship between the percentage fruit in diet and the size of the hippocampus or any of its subcomponents. While fruit acquisition may play an important role in intelligence [10, 86] and brain size [6], our findings suggest that the primary contribution of diet to these features may be the generalized support of the brain’s high metabolic costs [9] rather than specifically influencing neural systems specialized for spatial ability. On the other hand, non-dietary social-spatial memory factors, such as the ability to code for the locations of conspecifics, may be linked to hippocampus size.

Third, CA2 volume residuals showed no phylogenetic signal, except for in a regression with social and female group sizes in the single variable analysis (see Supplement for data on phylogenetic signal within each separate model). Thus, CA2 seems not to be under phylogenetic constraint related to home range or diet but is only shaped by social pressures. This finding can be interpreted as an indicator of the relative functional decoupling of this zone to the rest of the hippocampus. CA2 may show species-specific adaptations related to behavioural niche which deviate from trends within a clade. Recent work on the function of CA2 in mice found that it has a special role in social memory [66] and it has a different gene expression profile from CA1 and CA3 [47]. On the other hand, the adaptability of CA2 might come at a cost in terms of maintaining elementary functions shared across species - unlike CA1 and CA3 it is a smaller region and is not involved in spatial tasks [66]. Additionally, hilus was one of the structures that showed no relationship to social group size. It is important in spatial and memory functions and may be less adaptable to changes in social structure. We further investigated how hippocampus size is related to sexual dimorphism in primates since sex differences in hippocampal anatomy, spatial cognition, and home range size seem to be linked in some taxa [87]. We found that, of all brain structures examined, sexual dimorphism in the hippocampus is most closely related to somatic sexual dimorphism. It should be considered that spatial functions, like other brain functions, have become more corticalized in taxonomic groups with larger palliums such as primates [80]. However, the nature of the link is debated, for example, male superiority in spatial cognition may be a by-product of sex hormones rather than driven by ecological demands [88]. This provides a preliminary attempt to understand sex differences in the primate hippocampus.

Overall, we show that group size can predict the size of most hippocampus regions, while diet seems to be unrelated to hippocampal size at all. Moreover, group size was the only predictor that was related to total hippocampal size. Social group size is thought to be related to an increase in neocortex size, but this is mainly because of its role in higher cognitive social processes that are more demanding than simply remembering other individuals [89] [79]. Social memories seem to be structured within the spatial framework of the hippocampus too [25]. In fact, social memory might in part be an exaptation that “reuses” neural circuitry of the hippocampus for spatial maps in an ancestral mammal [90, 91]. In line with this, the role of hippocampus in spatial cognition is pronounced in rodents, but less well understood in primates; in humans, it is argued that the hippocampus appears to function in memory rather than spatial cognition [81]. Given the importance of social skills in primates, it is possible that in this order, social memory (overlain onto spatial maps originally for navigation) has increased in dominance over spatial mapping. The importance of the increasing evidence that social and spatial cognition rely on the same underlying representations in humans, such that spatial maps provide a means for mapping social relations, is developing into applications ranging from design considerations in the built environment to clinical implications [48].

We thank Anna van Oosterzee for stimulating discussions, Heiko Frahm for providing data and Simone Blomberg for statistical advice.

## Supporting information

Full results from all models

