## Supplementary material for "Primate hippocampus size and organization are predicted by sociality but not diet": data.html

| Species | Brainvol | Body.weight | Neo | Hippocampus.total | HP.HS.fibers | Hippocampus.retrocomm. | Subiculum | CA1 | CA2 | CA3 | Hilus | Fascia.dentata | Terrestriality | Diurnality | X.Fruit | Group.size | HR.size.avg | Diet\_cat\_dec | Soc\_sys\_dec | Mat\_sys\_dec | Soc\_gr\_sz\_dec | Group\_dunbar | GroupF\_dunbar |
| --- | --- | --- | --- | --- | --- | --- | --- | --- | --- | --- | --- | --- | --- | --- | --- | --- | --- | --- | --- | --- | --- | --- | --- |
|  |  |  |  |  |  |  |  |  |  |  |  |  |  |  |  |  |  |  |  |  |  |  |  |
| --- | --- | --- | --- | --- | --- | --- | --- | --- | --- | --- | --- | --- | --- | --- | --- | --- | --- | --- | --- | --- | --- | --- | --- |
| Alouatta\_seniculus | 49009 | 6400 | 31660 | 1319.79 | 315.03 | 1004.76 | 117.67 | 362.93 | 56.59 | 236.05 | 67.88 | 163.64 | 0 | 1 | 40.03333 | 7.872727 | 22.000 | Fol | Polygynandry | Harem Polygyny | 6.97 | 7.80 | 2.4 |
| Aotus\_trivirgatus | 16195 | 830 | 9950 | 538.80 | 133.49 | 405.31 | 27.43 | 150.16 | 20.83 | 97.14 | 33.52 | 76.23 | 0 | 0 | 65.00000 | 3.800000 | 10.000 | Frug | Pair | Monogamy | 3.51 | 2.90 | 1.0 |
| Ateles\_geoffroyi | 101034 | 8000 | 70856 | 1366.01 | 370.37 | 995.64 | 72.63 | 388.30 | 51.79 | 263.35 | 69.29 | 150.28 | 0 | 1 | 69.50000 | 31.000000 | 168.000 | Frug | Polygynandry | Polygynandry | 28.02 | 31.00 | 12.0 |
| Avahi\_laniger | 9798 | 1285 | 4813 | 526.25 | 67.70 | 458.55 | 48.13 | 178.89 | 15.29 | 112.49 | 24.93 | 78.82 | 0 | 0 | 0.00000 | 2.500000 | 1.500 | Fol | Pair | Monogamy | 2.67 | 2.50 | 1.0 |
| Avahi\_occidentalis | 9124 | 860 | 4443 | 475.58 | 56.15 | 419.43 | 41.61 | 175.41 | 9.13 | 90.39 | 30.86 | 72.03 | 0 | 0 | 0.00000 | 2.000000 | 0.700 | Fol | Pair | Monogamy | NA | 2.50 | 1.0 |
| Callicebus\_moloch | 17944 | 900 | 11163 | 588.29 | 101.92 | 486.37 | 42.71 | 174.88 | 22.90 | 129.11 | 37.33 | 79.44 | 0 | 1 | 54.00000 | 3.500000 | 9.250 | Frug | Pair | Monogamy | 3.37 | 3.50 | 1.0 |
| Callimico\_goeldii | 10510 | 480 | 6476 | 280.90 | 49.34 | 231.56 | 17.57 | 83.53 | 7.84 | 60.81 | 12.08 | 49.73 | 0 | 1 | 26.00000 | 4.500000 | 100.000 | Om | Pair | Monogamy | 6.50 | 8.00 | 2.0 |
| Callithrix\_jacchus | 7241 | 280 | 4371 | 221.12 | 40.63 | 180.49 | 11.48 | 73.07 | 7.49 | 42.82 | 12.29 | 33.34 | 0 | 1 | 20.60000 | 8.400000 | 4.600 | Frug | Pair | Polyandry | 7.88 | 9.50 | 4.0 |
| Callithrix\_pygmaea | 4302 | 120 | 2535 | 122.34 | 22.84 | 99.50 | 7.45 | 38.83 | 4.11 | 26.95 | 4.49 | 17.67 | 0 | 1 | 1.00000 | 5.400000 | 0.500 | Om | Pair | Polyandry | 5.63 | 5.63 | NA |
| Cebus\_albifrons | 66939 | 3100 | 46429 | 890.40 | 313.85 | 576.55 | 43.85 | 210.19 | 30.29 | 146.01 | 35.87 | 110.34 | 0 | 1 | NA | 25.000000 | 207.500 | Om | Polygynandry | Polygynandry | 21.13 | 19.80 | 7.3 |
| Cercopithecus\_ascanius | 63505 | 3400 | 45166 | 1189.00 | 249.60 | 939.40 | 77.57 | 362.05 | 23.02 | 240.60 | 85.70 | 150.46 | 0 | 1 | 40.20000 | 29.000000 | 34.000 | Frug | Polygyny | Harem Polygyny | 26.29 | 31.30 | 9.5 |
| Cercopithecus\_mitis | 70564 | 6300 | 49933 | 1366.39 | 379.53 | 986.86 | 77.48 | 378.84 | 30.44 | 288.47 | 86.05 | 125.58 | 0 | 1 | 50.00000 | 27.000000 | 48.500 | Frug | Polygynandry | Harem Polygyny | 21.28 | 20.70 | NA |
| Cheirogaleus\_major | 6373 | 450 | 2938 | 347.35 | 45.57 | 301.78 | 18.90 | 107.42 | 8.32 | 92.66 | 13.22 | 61.26 | 0 | 0 | NA | 4.000000 | 4.400 | Om | Solitary | Spatial Polygyny | 5.50 | 1.40 | NA |
| Cheirogaleus\_medius | 2961 | 117 | 1221 | 174.36 | 23.75 | 150.61 | 9.97 | 54.40 | 4.72 | 44.89 | 6.19 | 30.44 | 0 | 0 | 67.50000 | 4.000000 | 4.000 | Om | Solitary | Monogamy | 2.00 | 2.10 | 1.3 |
| Daubentonia\_madagascariensis | 42611 | 2800 | 22127 | 1776.10 | 328.20 | 1447.90 | 100.01 | 577.11 | 35.54 | 316.55 | 94.14 | 324.55 | 0 | 0 | 0.00000 | 1.000000 | 103.000 | Om | Solitary | Spatial Polygyny | 1.75 | 1.00 | NA |
| Erythrocebus\_patas | 103167 | 7800 | 77141 | 1590.53 | 504.92 | 1085.61 | 76.65 | 345.54 | 39.46 | 327.74 | 101.09 | 195.13 | 1 | 1 | 17.50000 | 25.000000 | 3200.000 | Om | Polygyny | Harem Polygyny | 26.52 | 34.80 | 14.0 |
| Eulemur\_fulvus\_fulvus | 22106 | 1400 | 12207 | 752.10 | 96.46 | 655.64 | 62.23 | 258.65 | 19.52 | 171.72 | 37.46 | 106.06 | 0 | 1 | 64.40000 | 11.500000 | 13.500 | Frug/Fol | Polygynandry | Polygynandry | 10.08 | 10.08 | NA |
| Galago\_senegalensis | 4512 | 186 | 2139 | 260.78 | 15.47 | 245.31 | 23.14 | 99.58 | 5.96 | 52.43 | 13.48 | 50.72 | 0 | 0 | NA | 3.500000 | NA | Om | Solitary | Spatial Polygyny | 1.00 | 2.50 | NA |
| Galagoides\_demidoff | 3203 | 81 | 1568 | 152.24 | 21.76 | 130.48 | 13.40 | 53.70 | 3.08 | 28.64 | 7.00 | 24.66 | 0 | 0 | 19.00000 | 4.000000 | 1.300 | Om | Solitary | Spatial Polygyny | 2.25 | 2.80 | NA |
| Gorilla\_gorilla\_gorilla | 470359 | 105000 | 341444 | 4780.48 | 1475.05 | 3305.43 | 191.70 | 1405.57 | 130.65 | 951.51 | 210.02 | 415.98 | 1 | 1 | 47.00000 | 10.500000 | 1770.000 | Fol | Polygyny | Harem Polygyny | 10.05 | 13.50 | NA |
| Homo\_sapiens | 1251847 | 65000 | 1006525 | 10287.38 | 2853.78 | 7433.60 | 821.76 | 2793.05 | 229.52 | 2357.63 | 397.17 | 834.47 | NA | NA | NA | NA | NA | NA | NA | NA | NA | NA | NA |
| Hylobates\_lar | 97505 | 5700 | 65800 | 2672.56 | 444.65 | 2227.91 | 139.39 | 866.51 | 76.73 | 547.00 | 226.25 | 372.03 | 0 | 1 | 63.70000 | 4.400000 | 48.000 | Frug | Pair | Monogamy | 3.51 | 4.50 | 1.0 |
| Indri\_indri | 36285 | 6250 | 20114 | 1529.38 | 358.58 | 1170.80 | 98.11 | 477.66 | 35.77 | 336.04 | 57.87 | 165.35 | 0 | 1 | 18.08500 | 4.000000 | 27.250 | Fol | Pair | Monogamy | 3.28 | 3.10 | 1.0 |
| Lagothrix\_lagotricha | 95503 | 5200 | 65873 | 1585.69 | 458.90 | 1126.79 | 78.34 | 400.11 | 49.57 | 332.56 | 73.93 | 192.28 | 0 | 1 | 66.70000 | 31.900000 | 700.000 | Om | Polygynandry | Polygynandry | 20.31 | 31.80 | 11.2 |
| Lepilemur\_ruficaudatus | 7175 | 915 | 3282 | 392.28 | 72.73 | 319.55 | 31.58 | 128.48 | 7.28 | 78.29 | 12.62 | 61.30 | 0 | 0 | NA | 1.500000 | 0.800 | Fol | Solitary | Spatial Polygyny | NA | 2.00 | 1.0 |
| Lophocebus\_albigena | 97603 | 7900 | 68733 | 1464.21 | 426.37 | 1037.84 | 73.47 | 395.87 | 38.15 | 304.20 | 69.90 | 156.25 | 0 | 1 | 41.00000 | 15.500000 | 225.000 | Frug | Polygynandry | Polygynandry | 15.68 | 16.20 | 7.0 |
| Loris\_tardigradus | 6269 | 322 | 3524 | 191.14 | 24.56 | 166.58 | 12.96 | 57.60 | 4.87 | 49.81 | 8.00 | 33.34 | 0 | 0 | 0.00000 | 2.500000 | 5.900 | Om | Solitary | Spatial Polygyny | 2.00 | 2.50 | 1.0 |
| Microcebus\_murinus | 1680 | 54 | 740 | 100.35 | 12.38 | 87.97 | 5.99 | 30.45 | 2.33 | 23.82 | 4.48 | 20.90 | 0 | 0 | 5.70000 | NA | 2.500 | Om | Solitary | Spatial Polygyny | 5.10 | 5.10 | NA |
| Miopithecus\_talapoin | 37776 | 1200 | 26427 | 704.87 | 190.75 | 514.12 | 41.31 | 183.40 | 15.31 | 134.60 | 51.91 | 87.59 | 0 | 1 | NA | NA | NA | Om | Polygynandry | Polygynandry | 68.38 | 64.00 | 27.0 |
| Nasalis\_larvatus | 92797 | 14000 | 62685 | 1965.93 | 518.83 | 1447.10 | 113.27 | 546.36 | 69.39 | 370.10 | 142.29 | 205.69 | 0 | 1 | 18.30000 | 12.700000 | 220.000 | Fol | Polygynandry | Harem Polygyny | 9.94 | 12.70 | 5.4 |
| Nycticebus\_coucang | 11755 | 800 | 6192 | 565.77 | 67.06 | 498.71 | 52.63 | 178.14 | 17.34 | 135.76 | 23.62 | 91.22 | 0 | 0 | 22.50000 | 2.000000 | 8.800 | Om | Solitary | Spatial Polygyny | 1.00 | 2.00 | 1.0 |
| Otolemur\_crassicaudatus | 9668 | 850 | 4723 | 460.88 | 73.43 | 387.45 | 42.22 | 164.58 | 9.60 | 78.22 | 28.14 | 64.69 | 0 | 0 | 21.00000 | 2.500000 | 8.500 | Om | Solitary | Spatial Polygyny | 2.25 | 2.40 | 1.0 |
| Pan\_troglodytes\_troglodytes | 382103 | 46000 | 291592 | 3778.69 | 1015.23 | 2763.46 | 266.58 | 1166.33 | 85.70 | 622.88 | 235.43 | 386.54 | 1 | 1 | 66.10000 | 40.700000 | 1983.333 | Frug | Polygynandry | Polygynandry | 42.71 | 42.71 | NA |
| Papio\_anubis | 190957 | 25000 | 140142 | 3397.90 | 1318.88 | 2079.02 | 156.72 | 800.97 | 66.37 | 564.35 | 178.36 | 312.25 | 1 | 1 | 38.00000 | 83.300000 | 2581.300 | Frug | Polygynandry | Polygynandry | 47.05 | 39.90 | 11.4 |
| Perodicticus\_potto | 13212 | 1150 | 6683 | 607.01 | 78.69 | 528.32 | 48.35 | 197.36 | 16.30 | 139.69 | 26.62 | 100.00 | 0 | 0 | 67.00000 | 2.000000 | 28.000 | Om | Solitary | Spatial Polygyny | 1.25 | 2.00 | 1.0 |
| Piliocolobus\_badius | 73818 | 7000 | 50906 | 1670.84 | 221.64 | 1449.20 | 129.20 | 615.79 | 38.44 | 323.72 | 156.77 | 185.28 | 0 | 1 | 24.10000 | 34.100000 | 55.500 | Fol | Polygynandry | Polygynandry | 29.18 | 34.10 | 13.2 |
| Pithecia\_irrorata | 32867 | 1500 | 21028 | 834.26 | 188.08 | 646.18 | 41.63 | 259.52 | 31.78 | 154.43 | 56.38 | 102.44 | 0 | 1 | 55.00000 | 3.750000 | 24.900 | NA | NA | NA | NA | 3.75 | NA |
| Propithecus\_verreauxi | 25194 | 3480 | 13170 | 1042.96 | 160.88 | 882.08 | 72.19 | 361.75 | 18.70 | 217.12 | 70.93 | 141.39 | 0 | 1 | 37.02400 | 5.500000 | 6.500 | Fol | Polygynandry | Polygynandry | 5.92 | 6.50 | 2.4 |
| Pygathrix\_nemaeus | 72530 | 7500 | 48763 | 2295.41 | 445.75 | 1849.66 | 117.14 | 786.60 | 44.48 | 420.73 | 227.64 | 253.07 | 0 | 1 | 25.50000 | 15.500000 | NA | Fol | Polygynandry | Harem Polygyny | 21.45 | 9.00 | 2.3 |
| Saguinus\_midas | 9569 | 340 | 5883 | 280.26 | 53.55 | 226.71 | 15.85 | 78.04 | 10.95 | 68.92 | 13.97 | 38.98 | 0 | 1 | 56.10000 | 5.900000 | 36.800 | Om | Pair | Polyandry | 5.00 | 4.00 | 1.0 |
| Saguinus\_oedipus | 9537 | 380 | 5894 | 262.09 | 53.10 | 208.99 | 23.45 | 85.42 | 9.66 | 49.77 | 9.14 | 31.55 | 0 | 1 | NA | 5.800000 | 13.900 | Om | Pair | Polyandry | 6.30 | 6.00 | NA |
| Saimiri\_sciureus | 22572 | 660 | 15541 | 351.44 | 103.06 | 248.38 | 17.58 | 79.82 | 13.70 | 66.43 | 21.07 | 49.78 | 0 | 1 | 31.00000 | 23.000000 | 97.500 | Om | Polygynandry | Polygynandry | 37.81 | 23.00 | 7.9 |
| Varecia\_variegata\_variegata | 29713 | 3000 | 15293 | 1403.65 | 226.70 | 1176.95 | 80.15 | 410.99 | 26.84 | 304.87 | 118.25 | 235.85 | 0 | 1 | 78.80000 | 5.800000 | 72.734 | Frug | Pair | Monogamy | 5.93 | 6.00 | NA |
