## Supplementary material for "Primate hippocampus size and organization are predicted by sociality but not diet": Social hippocampus R code.html

R Notebook


Code 

- Show All Code
- Hide All Code
- Download Rmd

### R Notebook

Try executing this chunk by clicking the *Run* button within the chunk or by placing your cursor inside it and pressing *Ctrl+Shift+Enter*.

Add a new chunk by clicking the *Insert Chunk* button on the toolbar or by pressing *Ctrl+Alt+I*.

When you save the notebook, an HTML file containing the code and output will be saved alongside it (click the *Preview* button or press *Ctrl+Shift+K* to preview the HTML file).

The preview shows you a rendered HTML copy of the contents of the editor. Consequently, unlike *Knit*, *Preview* does not run any R code chunks. Instead, the output of the chunk when it was last run in the editor is displayed.

### Load up packages


```
library(ape)
library(geiger)
library(nlme)
library(caper)
library(car)
```

### Preparation

#### Import data and tree


```
Data=read.csv("data_new.csv", header=TRUE)
TreePrimate=read.nexus("consensusTree_10kTrees_Primates_Version3.nex")
row.names(Data)=Data$Species
```

#### Convert into factor or numerical


```
Data$Brainvol <- as.numeric(Data$Brainvol)
Data$Body.weight<- as.numeric(Data$Body.weight)
Data$Neo<- as.numeric(Data$Neo)
Data$Hippocampus.total<- as.numeric(Data$Hippocampus.total)
Data$HP.HS.fibers<- as.numeric(Data$HP.HS.fibers)
Data$Hippocampus.retrocomm.<- as.numeric(Data$Hippocampus.retrocomm.)
Data$Subiculum<- as.numeric(Data$Subiculum)
Data$CA1<- as.numeric(Data$CA1)
Data$CA2<- as.numeric(Data$CA2)
Data$CA3<- as.numeric(Data$CA3)
Data$Hilus<- as.numeric(Data$Hilus)
Data$Fascia.dentata<- as.numeric(Data$Fascia.dentata)
Data$Terrestriality<- as.factor(Data$Terrestriality)
Data$Diurnality<- as.factor(Data$Diurnality)
Data$X.Fruit<- as.numeric(Data$X.Fruit)
Data$Group.size<- as.numeric(Data$Group.size)
Data$HR.size.avg<- as.numeric(Data$HR.size.avg)
Data$Soc_sys_dec<- as.factor(Data$Soc_sys_dec)
Data$Mat_sys_dec<- as.factor(Data$Mat_sys_dec)
Data$Soc_gr_sz_dec<- as.numeric(Data$Soc_gr_sz_dec)
Data$Group_dunbar<- as.numeric(Data$Group_dunbar)
Data$GroupF_dunbar<- as.numeric(Data$GroupF_dunbar)
```

#### Log and min-max stdize


```
Data$Brainvol <- log(Data$Brainvol)
Data$Body.weight<- log(Data$Body.weight)
Data$Neo<- log(Data$Neo)
Data$Hippocampus.total<- log(Data$Hippocampus.total)
Data$HP.HS.fibers<- log(Data$HP.HS.fibers)
Data$Hippocampus.retrocomm.<- log(Data$Hippocampus.retrocomm.)
Data$Subiculum<- log(Data$Subiculum)
Data$CA1<- log(Data$CA1)
Data$CA2<- log(Data$CA2)
Data$CA3<- log(Data$CA3)
Data$Hilus<- log(Data$Hilus)
Data$Fascia.dentata<- log(Data$Fascia.dentata)
Data$Group.size<- log(Data$Group.size)
Data$HR.size.avg<- log(Data$HR.size.avg)
Data$Soc_gr_sz_dec<- log(Data$Soc_gr_sz_dec)
Data$Group_dunbar<- log(Data$Group_dunbar)
Data$GroupF_dunbar<- log(Data$GroupF_dunbar)
#min-max stdze function
stdize = stdize = function(x, ...) {(x - min(x, ...)) / (max(x, ...) - min(x, ...))}
                        
Data$Brainvol <- stdize(Data$Brainvol, na.rm = T)
Data$Body.weight<- stdize(Data$Body.weight, na.rm = T)
Data$Neo<- stdize(Data$Neo, na.rm = T)
Data$Hippocampus.total<- stdize(Data$Hippocampus.total, na.rm = T)
Data$HP.HS.fibers<- stdize(Data$HP.HS.fibers, na.rm = T)
Data$Hippocampus.retrocomm.<- stdize(Data$Hippocampus.retrocomm., na.rm = T)
Data$Subiculum<- stdize(Data$Subiculum, na.rm = T)
Data$CA1<- stdize(Data$CA1, na.rm = T)
Data$CA2<- stdize(Data$CA2, na.rm = T)
Data$CA3<- stdize(Data$CA3, na.rm = T)
Data$Hilus<- stdize(Data$Hilus, na.rm = T)
Data$Fascia.dentata<- stdize(Data$Fascia.dentata, na.rm = T)
Data$X.Fruit<- stdize(Data$X.Fruit, na.rm = T)
Data$Group.size<- stdize(Data$Group.size, na.rm = T)
Data$HR.size.avg<- stdize(Data$HR.size.avg, na.rm = T)
Data$Soc_gr_sz_dec<- stdize(Data$Soc_gr_sz_dec, na.rm = T)
Data$Group_dunbar<- stdize(Data$Group_dunbar, na.rm = T)
Data$GroupF_dunbar<- stdize(Data$GroupF_dunbar, na.rm = T)
```

### plot all models


```
p <- ggplot(Data) + 
  theme_bw() +
  geom_jitter(aes(Hippocampus.total,Brainvol)) + geom_smooth(aes(Hippocampus.total,Brainvol), method=lm, colour = "black", se=FALSE) +
  geom_jitter(aes(Hippocampus.total, X.Fruit), colour="red", shape = 15) + geom_smooth(aes(Hippocampus.total, X.Fruit), colour = "red", method=lm, se=FALSE, linetype = "dashed") +
  geom_jitter(aes(Hippocampus.total, HR.size.avg), colour="blue", shape = 17) + geom_smooth(aes(Hippocampus.total, HR.size.avg), method=lm, colour="blue", se=FALSE, linetype = "longdash") +
  geom_jitter(aes(Hippocampus.total, Soc_gr_sz_dec), colour="darkgreen", shape = 3) + geom_smooth(aes(Hippocampus.total, Soc_gr_sz_dec), method=lm, colour="darkgreen", se=FALSE, linetype = "dotted") +
  labs(x = "ln(Hippocampus volume)", y = "ln(Total brain volume)    / Fraction Fruit    / ln(HR)    / Group Size     ")
```

### PGLS

### Create comp obj for caper


```
primate <- comparative.data(phy = TreePrimate, data = Data, names.col = Species, vcv = TRUE, na.omit = FALSE, warn.dropped = TRUE)
```


##### Neocortex


```
model.pglsNeo<-pgls(Hippocampus.total ~ Brainvol+Neo, data = primate, lambda='ML')
sink("neocortex.txt")
summary(model.pglsNeo)
anova(model.pglsNeo)
sink()
```

#### Home range


```
model.pgls1h<-pgls(Hippocampus.total ~ Brainvol+HR.size.avg, data = primate, lambda='ML')
model.pgls2h<-pgls(HP.HS.fibers ~ Brainvol+HR.size.avg, data = primate, lambda='ML')
model.pgls3h<-pgls(Hippocampus.retrocomm. ~ Brainvol+HR.size.avg, data = primate, lambda='ML')
model.pgls4h<-pgls(Subiculum ~ Brainvol+HR.size.avg, data = primate, lambda='ML')
model.pgls5h<-pgls(Hilus ~ Brainvol+HR.size.avg, data = primate, lambda='ML')
model.pgls6h<-pgls(CA1 ~ Brainvol+HR.size.avg, data = primate, lambda='ML')
model.pgls7h<-pgls(CA2 ~ Brainvol+HR.size.avg, data = primate, lambda='ML')
model.pgls8h<-pgls(CA3 ~ Brainvol+HR.size.avg, data = primate, lambda='ML')
model.pgls9h<-pgls(Fascia.dentata ~ Brainvol+HR.size.avg, data = primate, lambda='ML')
sink("model-HR.txt")
options(width=10000)
summary(model.pgls1h)
summary(model.pgls2h)
summary(model.pgls3h)
summary(model.pgls4h)
summary(model.pgls5h)
summary(model.pgls6h)
summary(model.pgls7h)
summary(model.pgls8h)
summary(model.pgls9h)
sink()
sink("model-HR-anova.txt")
options(width=10000)
anova(model.pgls1h)
anova(model.pgls2h)
anova(model.pgls3h)
anova(model.pgls4h)
anova(model.pgls5h)
anova(model.pgls6h)
anova(model.pgls7h)
anova(model.pgls8h)
anova(model.pgls9h)
sink()
```

#### Social models


```
model.pgls1s<-pgls(Hippocampus.total ~ Brainvol+Group.size, data = primate, lambda='ML')
model.pgls2s<-pgls(HP.HS.fibers ~ Brainvol+Group.size, data = primate, lambda='ML')
model.pgls3s<-pgls(Hippocampus.retrocomm. ~ Brainvol+Group.size, data = primate, lambda='ML')
model.pgls4s<-pgls(Subiculum ~ Brainvol+Group.size, data = primate, lambda='ML')
model.pgls5s<-pgls(Hilus ~ Brainvol+Group.size, data = primate, lambda='ML')
model.pgls6s<-pgls(CA1 ~ Brainvol+Group.size, data = primate, lambda='ML')
model.pgls7s<-pgls(CA2 ~ Brainvol+Group.size, data = primate, lambda='ML')
model.pgls8s<-pgls(CA3 ~ Brainvol+Group.size, data = primate, lambda='ML')
model.pgls9s<-pgls(Fascia.dentata ~ Brainvol+Group.size, data = primate, lambda='ML')
sink("model-group.txt")
options(width=10000)
summary(model.pgls1s)
summary(model.pgls2s)
summary(model.pgls3s)
summary(model.pgls4s)
summary(model.pgls5s)
summary(model.pgls6s)
summary(model.pgls7s)
summary(model.pgls8s)
summary(model.pgls9s)
sink()
sink("model-group-anova.txt")
options(width=10000)
anova(model.pgls1s)
anova(model.pgls2s)
anova(model.pgls3s)
anova(model.pgls4s)
anova(model.pgls5s)
anova(model.pgls6s)
anova(model.pgls7s)
anova(model.pgls8s)
anova(model.pgls9s)
sink()
####
model.pgls1sg<-pgls(Hippocampus.total ~ Brainvol+Soc_gr_sz_dec, data = primate, lambda='ML')
model.pgls2sg<-pgls(HP.HS.fibers ~ Brainvol+Soc_gr_sz_dec, data = primate, lambda='ML')
model.pgls3sg<-pgls(Hippocampus.retrocomm. ~ Brainvol+Soc_gr_sz_dec, data = primate, lambda='ML')
model.pgls4sg<-pgls(Subiculum ~ Brainvol+Soc_gr_sz_dec, data = primate, lambda='ML')
model.pgls5sg<-pgls(Hilus ~ Brainvol+Soc_gr_sz_dec, data = primate, lambda='ML')
model.pgls6sg<-pgls(CA1 ~ Brainvol+Soc_gr_sz_dec, data = primate, lambda='ML')
model.pgls7sg<-pgls(CA2 ~ Brainvol+Soc_gr_sz_dec, data = primate, lambda='ML')
model.pgls8sg<-pgls(CA3 ~ Brainvol+Soc_gr_sz_dec, data = primate, lambda='ML')
model.pgls9sg<-pgls(Fascia.dentata ~ Brainvol+Soc_gr_sz_dec, data = primate, lambda='ML')
sink("model-soc-group.txt")
options(width=10000)
summary(model.pgls1sg)
summary(model.pgls2sg)
summary(model.pgls3sg)
summary(model.pgls4sg)
summary(model.pgls5sg)
summary(model.pgls6sg)
summary(model.pgls7sg)
summary(model.pgls8sg)
summary(model.pgls9sg)
sink()
sink("model-soc-group-anova.txt")
options(width=10000)
anova(model.pgls1sg)
anova(model.pgls2sg)
anova(model.pgls3sg)
anova(model.pgls4sg)
anova(model.pgls5sg)
anova(model.pgls6sg)
anova(model.pgls7sg)
anova(model.pgls8sg)
anova(model.pgls9sg)
sink()
###
model.pgls1sd<-pgls(Hippocampus.total ~ Brainvol+Group_dunbar, data = primate, lambda='ML')
model.pgls2sd<-pgls(HP.HS.fibers ~ Brainvol+Group_dunbar, data = primate, lambda='ML')
model.pgls3sd<-pgls(Hippocampus.retrocomm. ~ Brainvol+Group_dunbar, data = primate, lambda='ML')
model.pgls4sd<-pgls(Subiculum ~ Brainvol+Group_dunbar, data = primate, lambda='ML')
model.pgls5sd<-pgls(Hilus ~ Brainvol+Group_dunbar, data = primate, lambda='ML')
model.pgls6sd<-pgls(CA1 ~ Brainvol+Group_dunbar, data = primate, lambda='ML')
model.pgls7sd<-pgls(CA2 ~ Brainvol+Group_dunbar, data = primate, lambda='ML')
model.pgls8sd<-pgls(CA3 ~ Brainvol+Group_dunbar, data = primate, lambda='ML')
model.pgls9sd<-pgls(Fascia.dentata ~ Brainvol+Group_dunbar, data = primate, lambda='ML')
sink("model-soc-group-dun.txt")
options(width=10000)
summary(model.pgls1sd)
summary(model.pgls2sd)
summary(model.pgls3sd)
summary(model.pgls4sd)
summary(model.pgls5sd)
summary(model.pgls6sd)
summary(model.pgls7sd)
summary(model.pgls8sd)
summary(model.pgls9sd)
sink()
sink("model-soc-group-dun-anova.txt")
options(width=10000)
anova(model.pgls1sd)
anova(model.pgls2sd)
anova(model.pgls3sd)
anova(model.pgls4sd)
anova(model.pgls5sd)
anova(model.pgls6sd)
anova(model.pgls7sd)
anova(model.pgls8sd)
anova(model.pgls9sd)
sink()
###
model.pgls1sdf<-pgls(Hippocampus.total ~ Brainvol+GroupF_dunbar, data = primate, lambda='ML')
model.pgls2sdf<-pgls(HP.HS.fibers ~ Brainvol+GroupF_dunbar, data = primate, lambda='ML')
model.pgls3sdf<-pgls(Hippocampus.retrocomm. ~ Brainvol+GroupF_dunbar, data = primate, lambda='ML')
model.pgls4sdf<-pgls(Subiculum ~ Brainvol+GroupF_dunbar, data = primate, lambda='ML')
model.pgls5sdf<-pgls(Hilus ~ Brainvol+GroupF_dunbar, data = primate, lambda='ML')
model.pgls6sdf<-pgls(CA1 ~ Brainvol+GroupF_dunbar, data = primate, lambda='ML')
model.pgls7sdf<-pgls(CA2 ~ Brainvol+GroupF_dunbar, data = primate, lambda='ML')
model.pgls8sdf<-pgls(CA3 ~ Brainvol+GroupF_dunbar, data = primate, lambda='ML')
model.pgls9sdf<-pgls(Fascia.dentata ~ Brainvol+GroupF_dunbar, data = primate, lambda='ML')
sink("model-soc-group-dunF.txt")
options(width=10000)
summary(model.pgls1sdf)
summary(model.pgls2sdf)
summary(model.pgls3sdf)
summary(model.pgls4sdf)
summary(model.pgls5sdf)
summary(model.pgls6sdf)
summary(model.pgls7sdf)
summary(model.pgls8sdf)
summary(model.pgls9sdf)
sink()
sink("model-soc-group-dunF-anova.txt")
options(width=10000)
anova(model.pgls1sdf)
anova(model.pgls2sdf)
anova(model.pgls3sdf)
anova(model.pgls4sdf)
anova(model.pgls5sdf)
anova(model.pgls6sdf)
anova(model.pgls7sdf)
anova(model.pgls8sdf)
anova(model.pgls9sdf)
sink()
```

#### Diet model


```
model.pgls1d<-pgls(Hippocampus.total ~ Brainvol+X.Fruit, data = primate, lambda='ML')
model.pgls2d<-pgls(HP.HS.fibers ~ Brainvol+X.Fruit, data = primate, lambda='ML')
model.pgls3d<-pgls(Hippocampus.retrocomm. ~ Brainvol+X.Fruit, data = primate, lambda='ML')
model.pgls4d<-pgls(Subiculum ~ Brainvol+X.Fruit, data = primate, lambda='ML')
model.pgls5d<-pgls(Hilus ~ Brainvol+X.Fruit, data = primate, lambda='ML')
model.pgls6d<-pgls(CA1 ~ Brainvol+X.Fruit, data = primate, lambda='ML')
model.pgls7d<-pgls(CA2 ~ Brainvol+X.Fruit, data = primate, lambda='ML')
model.pgls8d<-pgls(CA3 ~ Brainvol+X.Fruit, data = primate, lambda='ML')
model.pgls9d<-pgls(Fascia.dentata ~ Brainvol+X.Fruit, data = primate, lambda='ML')
sink("model-diet.txt")
options(width=10000)
summary(model.pgls1d)
summary(model.pgls2d)
summary(model.pgls3d)
summary(model.pgls4d)
summary(model.pgls5d)
summary(model.pgls6d)
summary(model.pgls7d)
summary(model.pgls8d)
summary(model.pgls9d)
sink()
sink("model-diet-anova.txt")
options(width=10000)
anova(model.pgls1d)
anova(model.pgls2d)
anova(model.pgls3d)
anova(model.pgls4d)
anova(model.pgls5d)
anova(model.pgls6d)
anova(model.pgls7d)
anova(model.pgls8d)
anova(model.pgls9d)
sink()
```

### Estimate ancestral states


```
#loading data
data=read.csv("data_new.csv", header=TRUE)
row.names(data)=data$Species
tree=read.nexus("consensusTree_10kTrees_Primates_Version3.nex")

#obraining residuals
primate <- comparative.data(phy = tree, data = data, names.col = Species, vcv = TRUE, na.omit = FALSE, warn.dropped = TRUE)
model.pgls.res<-pgls(log(Hippocampus.total) ~ log(Brainvol), data = primate, lambda='ML')

dat.rel <- residuals(model.pgls.res)
dat.abs <- as.matrix(log(data$Hippocampus.total), row.names=1)[,1]

#preparing data

dat.abs <- as.data.frame(dat.abs)
dat.rel <- as.data.frame(dat.rel)


rownames(dat.abs)<-data$Species
rownames(dat.rel)<-data$Species

names(dat.abs)[names(dat.abs) == 'dat.abs'] <- 'Absolute'
names(dat.rel)[names(dat.rel) == 'V1'] <- 'Relative'

dat.abs <- as.matrix(dat.abs)
dat.rel <- as.matrix(dat.rel)

dat.rel.vector <- as.vector(dat.rel)
names(dat.rel.vector) <- rownames(dat.rel)

dat.abs.vector <- as.vector(dat.abs)
names(dat.abs.vector) <- rownames(dat.abs)

#performing ANC

anc.abs <- fastAnc(tree, dat.abs, vars=TRUE,CI=TRUE, REML = 1)
anc.rel <- fastAnc(tree, dat.rel, vars=TRUE,CI=TRUE, REML = 1)

pdf(file = "ANC.pdf", width = 6, height = 6 , onefile = T)

fancyTree(tree,type="scattergram",
          X=as.matrix(dat.rel),A=as.matrix(anc.rel),control=list(spin=FALSE),label="horizontal")
fancyTree(tree,type="scattergram",
          X=as.matrix(dat.abs),A=as.matrix(anc.rel),control=list(spin=FALSE),label="horizontal")

dev.off ()
```

### Dimorphism


```
Data1=read.csv("dimorphdata.csv", header=TRUE)
row.names(Data1)=Data1$Species

TreeDimorph=read.tree("dimorphdata.txt")
plot(TreeDimorph, cex=0.5)

primate1 <- comparative.data(phy = TreeDimorph, data = Data1, names.col = Species, vcv = TRUE, na.omit = FALSE, warn.dropped = TRUE)

#sig
model.pgls1<-pgls(BoW ~ HIP, data = primate1, lambda='ML')
model.pgls2<-pgls(BoW ~ NET, data = primate1, lambda='ML')
model.pgls3<-pgls(BoW ~ OBL, data = primate1, lambda='ML')
model.pgls4<-pgls(BoW ~ CER, data = primate1, lambda='ML')
#sig
model.pgls5<-pgls(BoW ~  MES, data = primate1, lambda='ML')
model.pgls6<-pgls(BoW ~  DIE, data = primate1, lambda='ML')
model.pgls7<-pgls(BoW ~  TEL, data = primate1, lambda='ML')
model.pgls8<-pgls(BoW ~  BOL, data = primate1, lambda='ML')
#sig
model.pgls9<-pgls(BoW ~  PAL, data = primate1, lambda='ML')
model.pgls10<-pgls(BoW ~  SEP, data = primate1, lambda='ML')
model.pgls11<-pgls(BoW ~  STR, data = primate1, lambda='ML')
model.pgls12<-pgls(BoW ~  SCH, data = primate1, lambda='ML')
model.pgls13<-pgls(BoW ~  NEO, data = primate1, lambda='ML')
model.pgls14<-pgls(BoW ~  TOT, data = primate1, lambda='ML')

summary/anova
```
