## Supplementary material for "Primate hippocampus size and organization are predicted by sociality but not diet": supplementary.docx

**PGLS results from all four models including different measures of group size**

Table 1a. ANOVA output from testing the full model with (with Group size from Powell) vs hippocampal and regional volumes. Only predictors with p<0.0125 are marked with an asterisk. Degrees of freedom 29 and 4.

|  | Hippocampus | | HP+HS+ fibers | | Retrohippocampus | | Subiculum | | Hilus | | CA1 | | CA2 | | | CA3 | | | FD | |
| --- | --- | --- | --- | --- | --- | --- | --- | --- | --- | --- | --- | --- | --- | --- | --- | --- | --- | --- | --- | --- |
|  | Mean sq | p | Mean sq | p | Mean sq | p | Mean sq | p | Mean sq | p | Mean sq | p | Mean sq | p | Mean sq | | p | Mean sq | | p |
| Total Brain | 0.334 | <0.001* | 0.597 | <0.001* | 0.296 | <0.001* | 0.28 | <0.001* | 0.377 | <0.001* | 0.305 | <0.001* | 0.358 | <0.001* | 0.316 | | <0.001* | 0.243 | | <0.001* |
| Home Range | 0.002 | 0.1 | 0 | 0.95 | 0.004 | 0.04 | 0.121 | 0.02 | 0.008 | 0.05 | 0.011 | 0.003* | 0.003 | 0.09 | 0 | | 0.3 | 0 | | 0.6 |
| Group Size (Powell) | 0.009 | 0.003* | 0.004 | 0.06 | 0.011 | 0.002* | 0.006 | 0.07 | 0.007 | 0.07 | 0.011 | 0.003* | 0.011 | <0.001* | 0.012 | | 0.001* | 0.016 | | <0.001* |
| Fraction Fruit | 0.002 | 0.2 | 0 | 0.8 | 0.002 | 0.1 | 0 | 0.58 | 0.002 | 0.25 | 0.002 | 0.16 | 0.004 | 0.03 | 0.004 | | 0.04 | 0.002 | | 0.2 |
| Residuals | 0.024 |  | 0.027 |  | 0 |  | 0.001 |  | 0.002 |  | 0.001 |  | 0 |  | 0.001 | |  | 0.001 | |  |
| λ | 0.56 | | 0 | | 0.58 | | 0.49 | | 0.24 | | 0.57 | | 0 | | | 0.73 | | | 0.5 | |

Table 1b. ANOVA output from testing the full model with (with Social group size from DeCasien) vs hippocampal and regional volumes. Only predictors with p<0.0125 are marked with an asterisk. Degrees of freedom 28 and 4.

|  | Hippocampus | | HP+HS+ fibers | | Retrohippocampus | | Subiculum | | Hilus | | CA1 | | CA2 | | | CA3 | | | FD | |
| --- | --- | --- | --- | --- | --- | --- | --- | --- | --- | --- | --- | --- | --- | --- | --- | --- | --- | --- | --- | --- |
|  | Mean sq | p | Mean sq | p | Mean sq | p | Mean sq | p | Mean sq | p | Mean sq | p | Mean sq | p | Mean sq | | p | Mean sq | | p |
| Total Brain | 0.396 | <0.001* | 0.558 | <0.001* | 0.352 | <0.001* | 0.338 | <0.001* | 0.441 | <0.001* | 0.366 | <0.001* | 0.372 | <0.001* | 0.395 | | <0.001* | 0.282 | | <0.001* |
| Home Range | 0.002 | 0.09 | 0 | 0.68 | 0.005 | 0.022 | 0.013 | 0.008* | 0.006 | 0.08 | 0.011 | 0.002* | 0.004 | 0.04 | 0.001 | | 0.15 | 0 | | 0.61 |
| Social Group Size | 0.013 | <0.001* | 0.007 | 0.01* | 0.015 | <0.001* | 0.011 | 0.01* | 0.006 | 0.1 | 0.014 | <0.001* | 0.026 | <0.001* | 0.025 | | <0.001* | 0.018 | | <0.001* |
| Fraction Fruit | 0 | 0.44 | 0 | 0.93 | 0 | 0.33 | 0 | 0.76 | 0.001 | 0.39 | 0 | 0.41 | 0 | 0.39 | 0.002 | | 0.12 | 0 | | 0.50 |
| Residuals | 0 |  | 0.001 |  | 0 |  | 0 |  | 0.002 |  | 0.001 |  | 0.001 |  | 0 | |  | 0.001 | |  |
| λ | 0.62 | | 0.55 | | 0.61 | | 0.4 | | 0.23 | | 0.59 | | 0.63 | | | 0.84 | | | 0.55 | |

Table 1c. ANOVA output from testing the full model with (with Group size from Dunbar) vs hippocampal and regional volumes. Only predictors with p<0.0125 are marked with an asterisk. Degrees of freedom 30 and 4.

|  | Hippocampus | | HP+HS+ fibers | | Retrohippocampus | | Subiculum | | Hilus | | CA1 | | CA2 | | | CA3 | | | FD | |
| --- | --- | --- | --- | --- | --- | --- | --- | --- | --- | --- | --- | --- | --- | --- | --- | --- | --- | --- | --- | --- |
|  | Mean sq | p | Mean sq | p | Mean sq | p | Mean sq | p | Mean sq | p | Mean sq | p | Mean sq | p | Mean sq | | p | Mean sq | | p |
| Total Brain | 0.397 | <0.001* | 0.568 | <0.001* | 0.354 | <0.001* | 0.342 | <0.001* | 0.441 | <0.001* | 0.367 | <0.001* | 0.424 | <0.001* | 0.4 | | <0.001* | 0.280 | | <0.001* |
| Home Range | 0.002 | 0.074 | 0 | 0.58 | 0.005 | 0.02 | 0.014 | 0.007* | 0.009 | 0.04 | 0.012 | 0.001* | 0.003 | 0.05 | 0.001 | | 0.26 | 0 | | 0.57 |
| Group Size (Dunbar) | 0.012 | <0.001* | 0.011 | 0.002* | 0.013 | <0.001* | 0.008 | 0.04 | 0.007 | 0.06 | 0.013 | <0.001* | 0.014 | <0.001* | 0.019 | | <0.001* | 0.016 | | <0.001* |
| Fraction Fruit | 0.001 | 0.25 | 0 | 0.94 | 0.001 | 0.27 | 0 | 0.74 | 0.002 | 0.38 | 0 | 0.34 | 0.002 | 0.06 | 0.002 | | 0.09 | 0.001 | | 0.44 |
| Residuals | 0 |  | 0.001 |  | 0 |  | 0.001 |  | 0.002 |  | 0 |  | 0 |  | 0 | |  | 0.001 | |  |
| λ | 0.59 | | 0.56 | | 0.59 | | 0.47 | | 0.22 | | 0.58 | | 0 | | | 0.83 | | | 0.5 | |

Table 1d. ANOVA output from testing the full model with (with Female group size from Dunbar) vs hippocampal and regional volumes. Only predictors with p<0.0125 are marked with an asterisk. Degrees of freedom 20 and 4.

|  | Hippocampus | | HP+HS+ fibers | | Retrohippocampus | | Subiculum | | Hilus | | CA1 | | CA2 | | | CA3 | | | FD | |
| --- | --- | --- | --- | --- | --- | --- | --- | --- | --- | --- | --- | --- | --- | --- | --- | --- | --- | --- | --- | --- |
|  | Mean sq | p | Mean sq | p | Mean sq | p | Mean sq | p | Mean sq | p | Mean sq | p | Mean sq | p | Mean sq | | p | Mean sq | | p |
| Total Brain | 0.207 | <0.001* | 0.360 | <0.001* | 0.181 | <0.001* | 0.171 | <0.001* | 0.269 | <0.001* | 0.187 | <0.001* | 0.201 | <0.001* | 0.212 | | <0.001* | 0.135 | | <0.001* |
| Home Range | 0.004 | 0.013 | 0 | 0.7 | 0.009 | 0.003* | 0.0198 | 0.002* | 0.021 | 0.002* | 0.017 | <0.001* | 0.004 | 0.02 | 0.002 | | 0.14 | 0.004 | | 0.02 |
| Female Group Size | 0.008 | 0.002* | 0.005 | 0.03 | 0.009 | 0.003* | 0.009 | 0.02 | 0.003 | 0.09 | 0.007 | 0.02 | 0.009 | 0.002* | 0.014 | | <0.001* | 0.008 | | 0.002* |
| Fraction Fruit | 0 | 0.78 | 0 | 0.95 | 0 | 0.70 | 0 | 0.43 | 0 | 0.55 | 0.001 | 0.34 | 0.001 | 0.18 | 0 | | 0.3 | 0 | | 0.63 |
| Residuals | 0 |  | 0.001 |  | 0.001 |  | 0.001 |  | 0.001 |  | 0.001 |  | 0 |  | 0 | |  | 0.001 | |  |
| λ | 0.54 | | 0 | | 0.58 | | 0.39 | | 0 | | 0.6 | | 0 | | | 0.79 | | | 0.44 | |

**Testing separate models**

**(i) Ecological model: Home range**

We tested the prediction that variation in hippocampal and regional volumes can be predicted by variation in home range size. Only subiculum and CA1 were related to home range, after controlling for brain volume (subiculum: λ=0.57, F = 8.12, p=0.007 and CA1: λ=0.80, F = 9.02, p=0.005 on 2 and 36 degrees of freedom). See Table 2 and Supplementary for detailed results of all analysis.

**(ii) Ecological model: Diet**

We tested whether diet, that is, increase in frugivory, indicated as fraction fruit, can predict variation in hippocampal or regional volumes. We found no support for such hypothesis.

**(iii) Social model**

We tested whether variation in social group size can be predictive of variation in hippocampal or regional volumes and found support for such relation using 4 different measures of group size (see Methods - Social and ecological data). All measures of group size could explain volumetric variation in hippocampus and most subregions, except hilus (0 out of 4 measures), HP+HS+fibers (1 out of 4), and subiculum (3 out of 4). Interestingly, CA2 volume showed no phylogenetic signal in any ecological or social model, except for when tested against social group size and female group size (see Table 3 for phylogenetic signal of the residuals in every model).

**Table 2.** Sequential SS for PGLS results of home range, fraction fruit, group size (Powell) (S1), social group size (S2), group size (Dunbar) (S3), and female group size (S4) controlling for total brain volume. DF - degrees of freedom for each model. For list of abbreviations, see Methods – Anatomical data and Table 1.

Only predictors with p<0.0083 are filled in grey. See supplement for full test results, including F, t statistics and exact p values.

|  | HIP | HP+HS+ fibers | RH | Subiculum | Hilus | CA1 | CA2 | CA3 | FD | DF |
| --- | --- | --- | --- | --- | --- | --- | --- | --- | --- | --- |
| Total brain |  |  |  |  |  |  |  |  |  |  |
| Home Range |  |  |  |  |  |  |  |  |  | 36 |
| Fraction fruit |  |  |  |  |  |  |  |  |  | 33 |
| S1 |  |  |  |  |  |  |  |  |  | 37 |
| S2 |  |  |  |  |  |  |  |  |  | 36 |
| S3 |  |  |  |  |  |  |  |  |  | 39 |
| S4 |  |  |  |  |  |  |  |  |  | 26 |

**Table 3.** Phylogenetic signal (λ) of the residuals in the regression models for each hippocampal component tested from Table 2. All models include total brain size. Group size (Powell) (S1), social group size (S2), group size (Dunbar) (S3), and female group size (S4).

|  | HIP | HP+HS+ fibers | RH | Subiculum | Hilus | CA1 | CA2 | CA3 | FD |
| --- | --- | --- | --- | --- | --- | --- | --- | --- | --- |
| λ Home Range | .67 | .23 | .77 | .57 | .27 | .80 | 0 | .74 | .79 |
| λ Fraction Fruit | .80 | .06 | .99 | .80 | .64 | .98 | 0 | .86 | .81 |
| λ S1 | .60 | .21 | .64 | .53 | .49 | .62 | 0 | .72 | .50 |
| λ S2 | .64 | .23 | .66 | .42 | .52 | .62 | .84 | .78 | .55 |
| λ S3 | .68 | .34 | .73 | .59 | .63 | .71 | .28 | .85 | .51 |
| λ S4 | .67 | .12 | .8 | .68 | .73 | .75 | 0 | .86 | .59 |
