## Supplementary figures and images for "Primate hippocampus size and organization are predicted by sociality but not diet"

### Summary of missing data.pdf

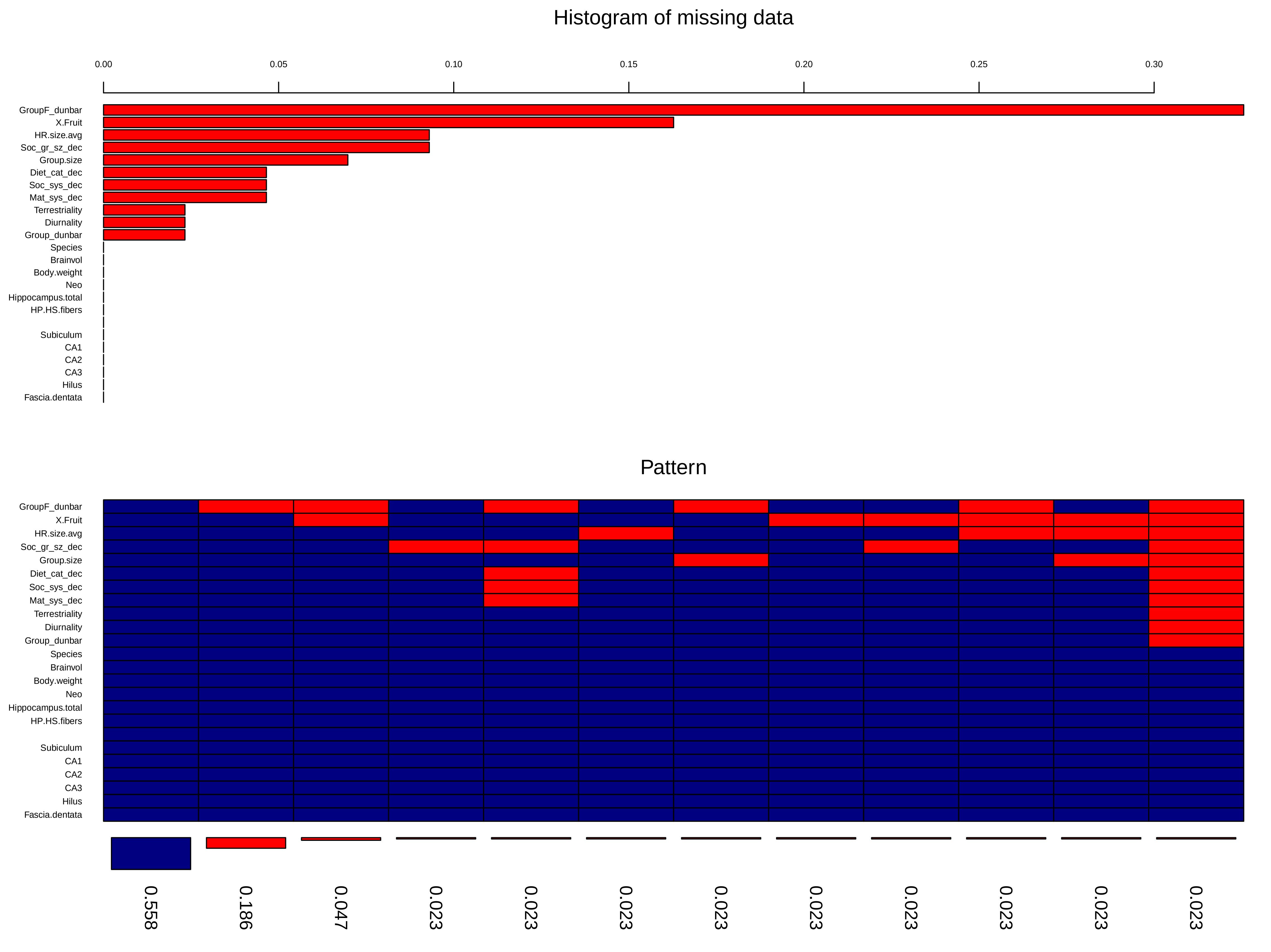
